# DIRECT DETECTION OF *LISTERIA MONOCYTOGENES* BY RECOMBINASE POLYMERASE AMPLIFICATION

**DOI:** 10.1101/2021.05.22.445288

**Authors:** Hau Thi Tran, Diem Hong Tran, Trang Nguyen Minh Pham, Huong Thi Thu Phung

**Affiliations:** NTT Hi-Tech Institute, Nguyen Tat Thanh University, 298A Nguyen Tat Thanh, Ho Chi Minh City, 700000, Vietnam.

**Keywords:** *Listeria monocytogenes*, direct RPA, foodborne diseases, rapid detection, isothermal PCR

## Abstract

*Listeria monocytogenes* is one of the most common types of food poisoning bacteria which can cause serious foodborne diseases or even lethality. Generally, *L. monocytogenes* can be detected using traditional microbiology or molecular biology techniques, notably PCR. However, the application of these methods at the field is restricted due to the strict requirement of equipment and skilled personnel. In this study, recombinase polymerase amplification (RPA), an isothermal PCR assay was developed to rapidly detect *L. monocytogenes* in the crude samples. The results showed that the RPA reaction, without requiring complex thermal cycles, was well-performed in the optimal conditions of 39°C within only 25 minutes. The limit of detection was identified as 310 fg of *L. monocytogenes* genomic DNA, which was 1000-fold more sensitive than the conventional PCR. In addition, RPA also succeeded to directly detect *L. monocytogenes* cells at a concentration as low as 2.5 × 10^1^ Colony Forming Unit (CFU)/ml in pure cultures and 2.5 × 10^2^ CFU/ml in crude samples without sample extraction or processing. Therefore, RPA established in this study could be an alternative standard method to confirm the presence of *L. monocytogenes* in food. Accordingly, this rapid and sensitive method could be further applied to clinical testing for the diagnosis of *L. monocytogenes* infection, especially in areas with limited settings.

## INTRODUCTION

*Listeria monocytogenes* is a Gram-positive, rod-shaped, facultatively anaerobic and non-sporulating bacterium, which is the causative agent of human listeriosis, a rare foodborne infectious disease with high hospitalization and case lethality rates (**Gandhi & Chikindas, 2007; Carvalho *et al.*, 2014**). The major sources of *L. monocytogenes* infection were unpasteurized milk or soft cheese made from raw milk. Besides, consumption of the ready-to-use meat contaminated was also considered as an important risk source of the *L. monocytogenes* infection (**Swaminathan & Gerner-Smidt, 2007**). In 2018, an outbreak of listeriosis was reported with 978 confirmed cases in South Africa. Most of the cases are neonates, pregnant women, the elderly, and immunocompromised persons (**WHO, 2018**). In Vietnam, the serious consequences of human listeriosis infection causing meningitis were reported as well (**Chau *et al.*, 2010**).

Commonly, *L. monocytogenes* can be detected using traditional microbiology or biomolecular techniques. The traditional microbiological methods for detecting *L. monocytogenes* are quite time-consuming and sophisticated (**Curtis & Lee, 1995**). A suitable culture media is required for the growth of *L. monocytogenes* and an identified step was performed with some complex biochemical assays before isolating *L. monocytogenes* from cultures (**Taherkhani *et al.*, 2013**). In recent decades, PCR was considered as an effective molecular method to alternate the conventional microbiological method to detect different bacterial pathogens including *L. monocytogenes* (**Göksoy & Kaya, 2006; Swetha *et al.*, 2016**). The method has significant improvements in sensitivity and specificity. However, PCR requires an accurate thermal cycler and a considerable running time while the application of the standard operating procedure for PCR at point-of-care diagnosis is restricted (**Delibato *et al.*, 2009**).

Recently, isothermal amplification approaches based on conventional PCR assay such as loop-mediated isothermal amplification (LAMP), cross-priming amplification (CPA), polymerase spiral reaction (PSR) and recombinase polymerase amplification (RPA) have been widely used in analyzing the foodborne organism pathogens (**Notomi *et al.*, 2000; Craw & Balachandran, 2012; Xu *et al.*, 2012; Craw *et al.*, 2013; W. Liu *et al.*, 2015**). LAMP, CPA and PSR assays were previously employed to detect *L. monocytogenes* (**Wang *et al.*, 2014; Du *et al.*, 2018; Nathaniel *et al.*, 2019**). All of these methods were shown to have high sensitivity and specificity with the ideal limit of detection (LOD). Moreover, these assays are simpler, more rapid and cost-effective compared to conventional PCR.

LAMP, CPA, and PSR have been considered useful isothermal amplification techniques for the early diagnosis of *L. monocytogenes*. However, these methods usually require sets of specially designed primers for identifying distinct regions of the target sequence as well as higher incubation temperatures (**Piepenburg *et al.*, 2006**). The RPA assay possesses some advantages over the others because it requires only two specific primers and a lower incubation temperature to run the reaction. The amplification of nucleic acid in RPA is performed by a recombinase-primer complex. A DNA polymerase having a strand-displacement activity is utilized to extend the specific primers at cognate sites and the intermediate structures are stabilized by single-stranded DNA binding proteins. Additionally, the RPA reactions do not need a global melting step of the template, thus, the requirements of restricted equipment for RPA assay are not essential (**Piepenburg *et al.*, 2006**). Nowadays, portable and compact lateral flow (LF) strips have already provided the ideal method for simple and rapid visualization of the RPA amplicon at the field (**Daher *et al.*, 2016**). LF strips utilize hybridisation products as the substrate, for example RPA products and antigen-labelled probes. LF strips are labelled with an antibody specific to an antigen labelled on probes accordingly. Typically, amplicon can be detected by capture of antigen by anti-FAM and anti-Biotin antibodies, displaying a visual band on the strips (**Daher *et al.*, 2016**). Additionally, RPA amplified product can be applied directly onto the dipstick strips without purification, generating results within merely 5 min afterward (**Daher *et al.*, 2016**). Thus, the result can be read by the naked eye shortly. Therefore, combination with LF strips can minimize and simplize the detection procedure of RPA product in a resource-limited setting.

Previously, the RPA assays were successfully established to identify some types of food poisoning bacteria (**Gao *et al.*, 2017; Liu *et al.*, 2017; Du *et al.*, 2018; Geng *et al.*, 2019**; **Hu *et al.*, 2020**). Previously, **Gao** *et al.* (**2017**) utilized RPA to detect *L. monocytogenes* successfully with a limit of detection (LOD) of 10 pg of genomic DNA per reaction. However, they did not perform direct detection of *L. monocytogenes* cells in simulated samples. Later, **Du** *et al.* (**2018**) evaluated the RPA performance for detecting *L. monocytogenes* with the LOD of 300 fg of genomic DNA and 1.5 Colony Forming Unit (CFU) in spiked food samples. Nevertheless, their approach required sample enrichment for 6 hours. Therefore, in this study, we attempted to establish a direct RPA assay for rapid and accurate detection of *L. monocytogenes* cells in the unprocessed food sample. The simply operational mechanism, the isothermal establishment and the short testing time make the RPA assay developed more accessible to limited setting areas.

## MATERIAL AND METHODS

### Bacterial cultivation

*L. monocytogenes* (laboratory collection) was cultured overnight at 37°C in Brain Heart Infusion broth (HiMedia Laboratories Pvt. Ltd, India). Through shaking bacterial cultures at 180 rounds per min (rpm), precipitation of cells was avoided. Other bacterial strains (laboratory collection) including *Salmonella enterica, Staphylococcus aureus, Clostridium perfringens, Bacillus cereus,* and *Vibrio parahaemolyticus* were cultured similarly.

### DNA extraction

The cetrimonium bromide (CTAB) extraction buffer contains 2% (w/v) CTAB, 100 mM Tris-HCl (pH 8), 20 mM EDTA (pH 8), 1.4 M NaCl. The *L. monocytogenes* cells were harvested from 1 ml of culture by centrifugation, and supernatants were then discarded. Cells were resuspended in 800 μl of the pre-warmed (65°C) CTAB lysis buffer and mixed thoroughly, then incubated at 65°C for 60 min. Samples were then centrifuged at 4°C for 15 min at 14000 *g*. Supernatants were transferred to fresh tubes and an approximately equal volume of Phenol: Chloroform: Isoamyl alcohol (PCI) was added and mixed thoroughly. Phase separation occurred by centrifugation at 14000 *g* for 15 min at 4°C. The upper aqueous phases were moved to new tubes and a 2.5 equal volume of ethanol 99% was added. DNA was precipitated overnight at −20°C. The DNA was then dissolved with 50 μl of elution buffer. The concentration and purification of DNA were measured with a Genova Plus Spectrophotometer (Jenway, Staffordshire, UK). The extracted genomic DNA was then stored at −20°C until use.

### Primer design

The RPA primers targeting the *plcA* gene of *L. monocytogenes* (Gene bank: CP023861.1) were designed according to the guidelines provided by TwistDx (Cambridge, UK) (http://hdl.handle.net/20.500.11794/26269). The primer set was chosen by assessing the specificity using NCBI BLAST and the amplification effects were also evaluated practically. *In silico* PCR analysis function available in FastPCR software (http://primerdigital.com/fastpcr.html) was additionally utilized for primer specificity analysis. The primer sequences were aligned to the reference genome sequences of other strains downloaded from NCBI. Two candidate primer pairs were commercially synthesized by IDT (Singapore). The primer pairs that produced the clearly visible bands representing for amplified product in agarose gel electrophoresis were selected and the sequences were listed in Table 1.

**Table 1.**
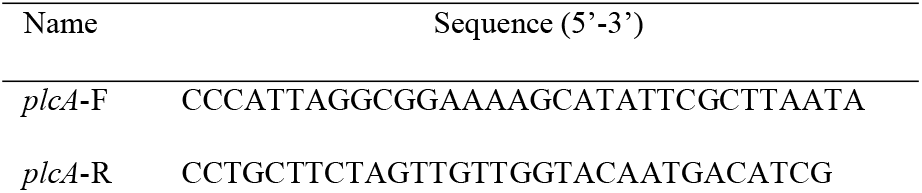
Primer sequence for *L. monocytogenes* RPA assay

### PCR reaction

The PCR was carried out in a reaction volume of 20 μl in small tubes containing 0.4 μM each of RPA primers, 4 μl of 5X Mytaq reaction buffer (Bioline, London, UK), 0.2 μl of MyTaq DNA polymerase (Bioline, London, UK) and 1 μl of DNA template. The PCR reaction was run as follows: initial denaturation stage at 95°C for 2 min, 35 cycles of 95°C for 30 seconds, 58°C for 30 seconds, 72°C for 30 seconds, and final extension stage at 72°C for 10 min. The PCR results were analyzed by a 1.5% agarose gel electrophoresis.

### RPA reaction for detection of *L. monocytogenes*

The RPA assay was performed referring to the TwistDx’s recommended protocols (http://hdl.handle.net/20.500.11794/26269). The reaction mixture containing 29.5 μl of a rehydration buffer, 2.4 μl of 10 μM forward and reverse primer, 12.2 μl of sterile water and 1 μl of the template was transferred to a lyophilized pellet tube. Then, 2.5 μl of magnesium acetate was added to start the reaction. Sterile water was used as a negative control sample. The tubes were mixed thoroughly and then centrifuged briefly. Subsequently, the tubes were incubated at 39°C for 25 min in BioSan Dry block thermostat Bio TDB-100. A mixing step after 4 min of incubation was carried out for better sensitivity of the assay. Finally, 50 μl of PCI was added to the tubes and vortexed lightly. The tubes were centrifuged at 14000 *g* for 10 min to remove the undesirable components affecting the read-out of the results. The RPA products were analyzed by a 1.5% agarose gel electrophoresis and visualized under the UV light using G: BOX Mini 6/9 (Syngene, Cambridge, UK).

### Optimization of RPA reaction

According to the manufacturer’s guidelines, the effective incubation temperature of TwistAmp Basic kit (https://www.twistdx.co.uk/en/products/product/twistamp-basic) used in this study ranges between 37°C - 39°C. Therefore, to determine the optimal reaction temperature and incubation time of the RPA reaction, the RPA assays were performed at temperatures ranging from 35°C to 41°C for different times including 15, 20, 25 and 30 min. The amount of 31 pg of genomic DNA was used as the template for optimizing the *L. monocytogenes* RPA assay.

### Evaluation of Specificity of RPA assay

The specificity of the RPA reaction was assessed under the optimal temperature and incubation time determined. Cross-reactivity analysis using the extracted genomic-DNAs of other typical foodborne pathogens including *S. enterica, S. aureus, C. perfringens, B. cereus,* and *V. parahaemolyticus* (laboratory collection) was also performed.

### Evaluation of LOD of RPA assay

To evaluate the LOD of the RPA assay, a 10-fold serial dilution from 310 ng to 3.1 fg of the extracted genomic-DNA of *L. monocytogenes* was prepared. One μl of each DNA concentration was utilized as the template for RPA and PCR assays. The LOD of the RPA reaction was compared with the LOD of the PCR assay.

To determine the LOD of the direct RPA assay, various concentrations of the *L. monocytogenes* cell culture were prepared. *L. monocytogenes* was initially cultured in 10 ml of fresh Brain Heart Infusion broth at 37°C for 24 hours. The *L. monocytogenes* cell concentration was determined using the count plating method. Next, 1 ml of the *L. monocytogenes* culture was centrifuged at 4000 *g* for 20 min at 4°C to harvest the cells. The final pellet was washed two times and resuspended in 100 μl of 0.9% NaCl. A serial dilution of the *L. monocytogenes* cells was then prepared to attain samples with a final concentration ranging from 2.5 × 10^8^ to 2.5 × 10^0^ CFU/ml. Then one μl of each cell concentration was directly utilized as the template for the RPA assay.

### Direct detection of *L. monocytogenes* in contaminated milk

To evaluate the efficiency of RPA assay for the direct detection of *L. monocytogenes* in contaminated milk, the cell pellets harvested from different concentrations of cell cultures were spiked into 1 ml of the pasteurized milk purchased from the local supermarket. The tubes were vortexed and centrifuged at 6000 *g* for 3 minutes. Next, 900 μl of the upper liquid was removed gently. Then, each sample was used as the template for direct RPA assays.

## RESULTS AND DISCUSSION

### RPA assay for detection of *L. monocytogenes*

RPA primers were designed to detect the *plcA* gene of *L. monocytogenes*. The target region is an important virulence gene that has been shown to have high specificity for diagnosing *L. monocytogenes* strain (**Lida *et al.*, 2014**). The RPA reactions were performed using 3.1 ng of the genomic DNA of *L. monocytogenes* as a template. The result showed that the assay produced a clearly visible band at approximately 228 base pairs (bp) of the expected size when analyzed by gel electrophoresis (Fig. 1, lane 4). The size of the amplified product obtained by RPA is similar to the amplicon produced by PCR when using the same primer set (Fig. 1, lane 2), indicating that the RPA primers designed successfully amplified the target sequence as expected.

**Figure 1.**
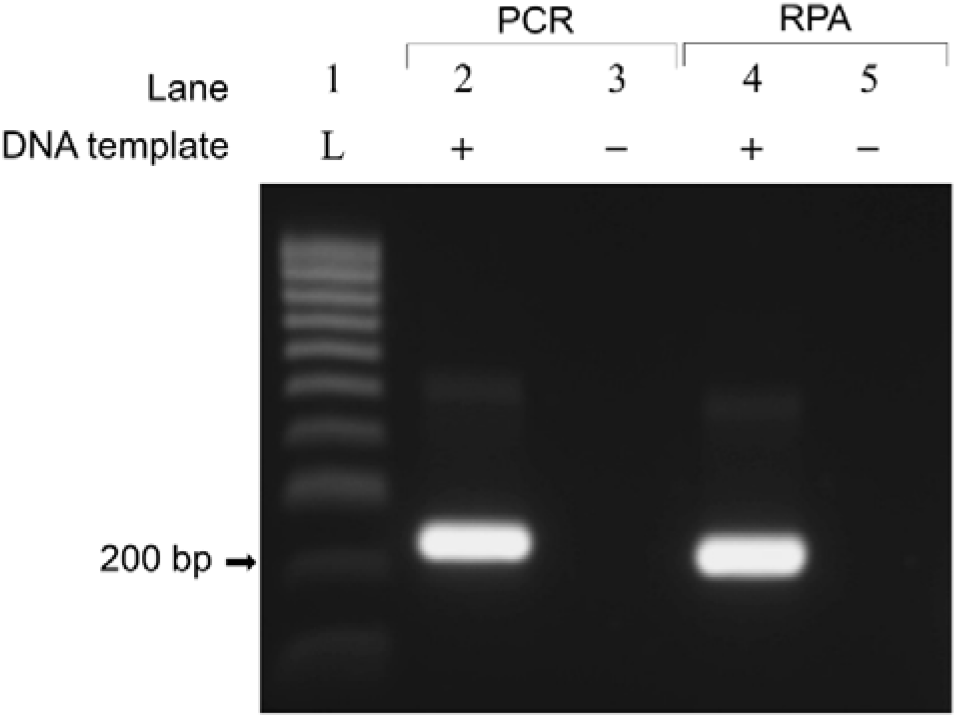
Detection of genomic DNA of *L. monocytogenes* by RPA and PCR. Positive reactions contain 3.1 ng of the DNA template. Sterile water is used as the negative control sample. Abbreviation, L: DNA ladder.

### Optimization of RPA assay for detection of *L. monocytogenes*

The optimal temperature and incubation time of the RPA reaction for the detection of *L. monocytogenes* were determined. The results indicated that the highest amount of amplified product was observed at 39°C (Fig. 2A, lane 4). For incubation time, the RPA amplicon could be seen just after 15 min and got saturation after 25 min (Fig. 2B). Thus, the optimal condition of the RPA reaction for detection of *L. monocytogenes* genomic DNA was set at 39°C for 25 min.

**Figure 2.**
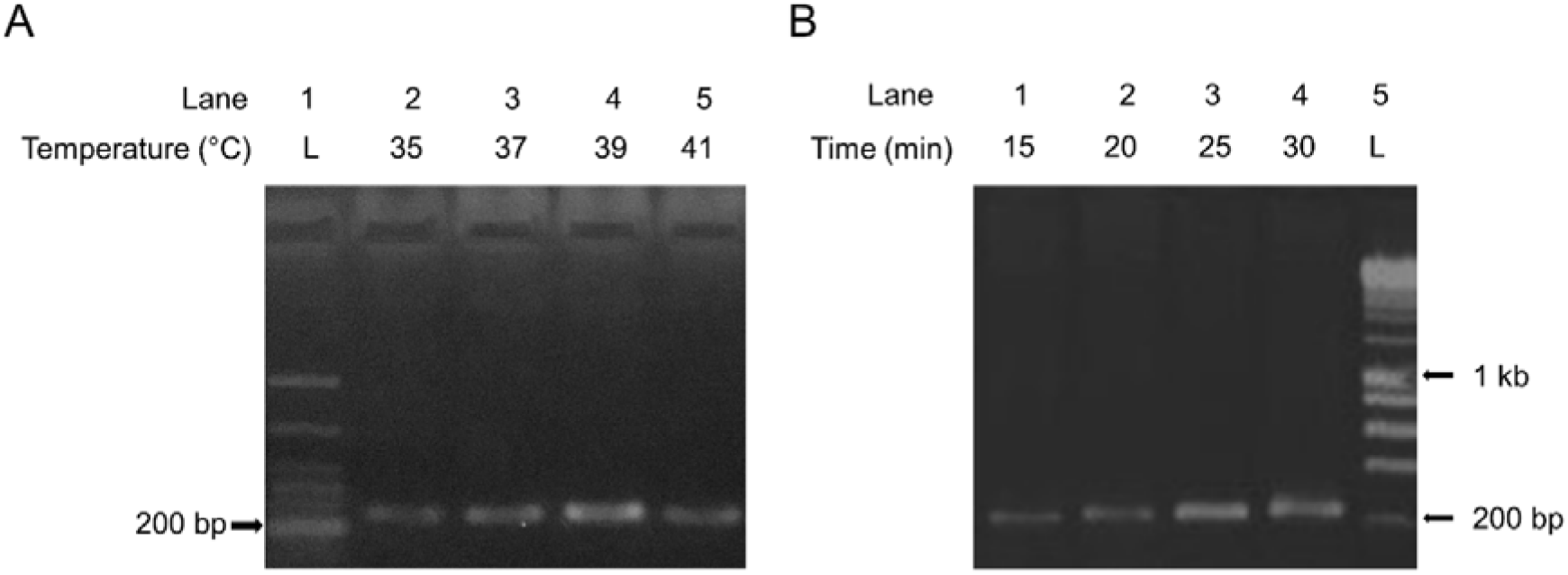
Optimization of the temperature and incubation time of RPA reaction. The reaction contains 31 pg of *L. monocytogenes* genomic DNA. (A) The RPA reactions were incubated at 35, 37, 39, and 41°C, respectively. (B) The RPA reactions were incubated at 39°C from 15 to 30 min. Abbreviation, L: DNA ladder.

### Specificity of RPA assay for detection of *L. monocytogenes*

*In silico* PCR analysis showed that the designed primer pair would not amplify the genome sequences of 20 different bacterial strains, supporting that the selected primer set has high specificity for identifying *L. monocytogenes* (Table 2). To practically evaluate the specificity of RPA assay developed for detection of *L. monocytogenes*, genomic DNAs of several bacteria commonly causing food poison were extracted and used as the template for RPA reactions. The results indicated that no cross-reactivity was observed with the foodborne bacterial strains examined including *S. enterica, S. aureus, C. perfringens, B. cereus,* and *V. parahaemolyticus* (Fig. 3). Thus, the RPA primer pair designed is highly specific for *L. monocytogenes*.

**Table 2.** *In silico* PCR results (0-2 mismatch allowed in 3’-end)

| No | Organism | Primer binding | PCR product |
| --- | --- | --- | --- |
| 1 | *L. monocytogenes* | + | + |
| 2 | *Bacillus anthracis* | – | – |
| 3 | *B. cereus* | – | – |
| 4 | *Campylobacter jejuni* | – | – |
| 5 | *Candida albicans* | – | – |
| 6 | *Clostridium botulinum* | – | – |
| 7 | *C. perfringens* | – | – |
| 8 | *Escherichia coli* | – | – |
| 9 | *Leptospira interrogans* | – | – |
| 10 | *Mycobacterium tuberculosis* | – | – |
| 11 | *Neisseria meningitidis* | – | – |
| 12 | *Pseudomonas aeruginosa* | – | – |
| 13 | *S. enterica* | – | – |
| 14 | *Shigella sonnei* | – | – |
| 15 | *S. aureus* | – | – |
| 16 | *Staphylococcus epidermidis* | – | – |
| 17 | *Streptococcus pneumoniae* | – | – |
| 18 | *Streptococcus salivarius* | – | – |
| 19 | *Vibrio cholerae* | – | – |
| 20 | *V. parahaemolyticus* | – | – |
| 21 | *Yersinia pseudotuberculosis* | – | – |

**Figure 3.**
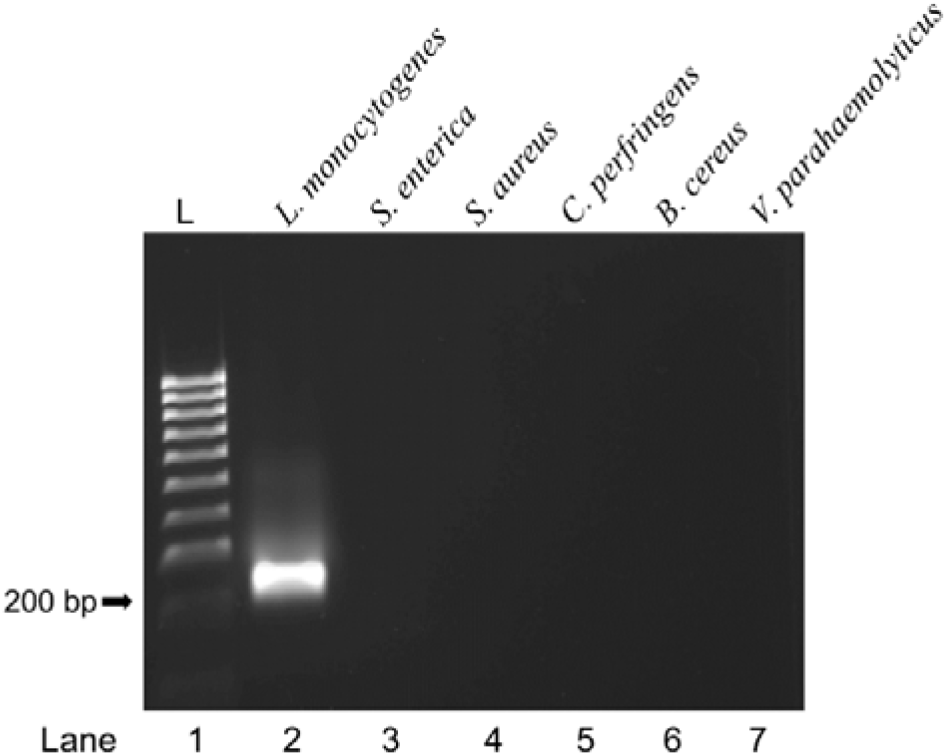
The specificity of *L. monocytogenes* RPA assay was evaluated with 3.1 ng of genomic DNAs of *L. monocytogenes* and other foodborne bacteria. Abbreviation, L: DNA ladder.

### Detection limit of RPA assay for detection of *L. monocytogenes*

Evaluation of the sensitivity of *L. monocytogenes* RPA assay developed was performed using a ten-fold dilution of extracted DNA of *L. monocytogenes*. The results showed that the lowest amount of extracted-genomic DNA that RPA could detect was 310 fg/reaction which equivalents to 99 genome copies per reaction (Fig. 4A). Meanwhile, the LOD value of PCR utilizing the same primer set was identified at 310 pg/reaction (Fig. 4B). The RPA assay is thus approximately 1000 times more sensitive than the PCR reaction in this study.

**Figure 4.**
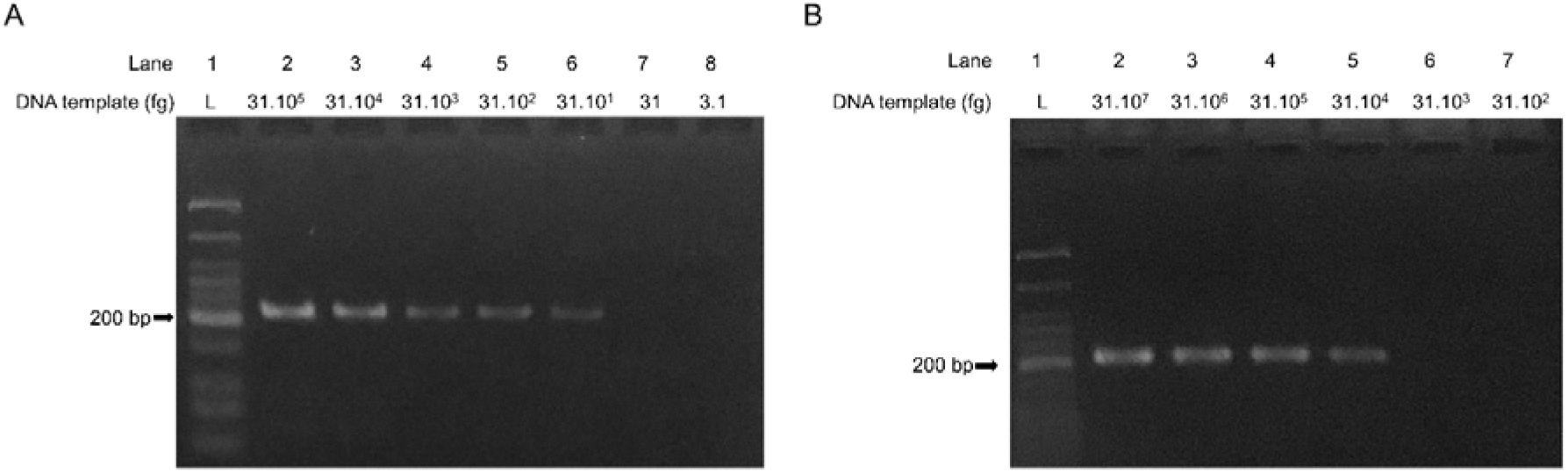
Comparison of the LOD values between RPA and PCR assay for detection of *L. monocytogenes* genomic DNA. The LOD values of RPA (A) and PCR (B) were evaluated using the ten-fold serial dilution of *L. monocytogenes* genomic DNA extracted ranging from 310 ng to 3.1 fg. One μl of each DNA concentration was used for the PCR and RPA reactions, respectively. Abbreviation, L: DNA ladder.

### Direct RPA assay for detection of *L. monocytogenes*

We attempted to detect *L. monocytogenes* cells from unextracted samples by the RPA assay developed. To this end, the cell culture of *L. monocytogenes* was used directly as the template for RPA reactions. As expected, the RPA assay could detect the presence of *L. monocytogenes* genomic DNA without the sample extraction process (Fig. 5A). Next, the LOD of direct RPA for detection of *L. monocytogenes* was determined using the serial dilution of *L. monocytogenes* cell culture. The results showed that the amplified products could be observed at the cell concentrations ranging from 2.5 × 10^6^ to 2.5 × 10^1^ CFU/ml (Fig. 5B). There was no RPA amplicon produced at 2.5 CFU/ml (Fig. 5B, lane 7). Thus, the LOD of *L. monocytogenes* direct RPA was determined at 2.5 × 10^1^ CFU/ml.

**Figure 5.**
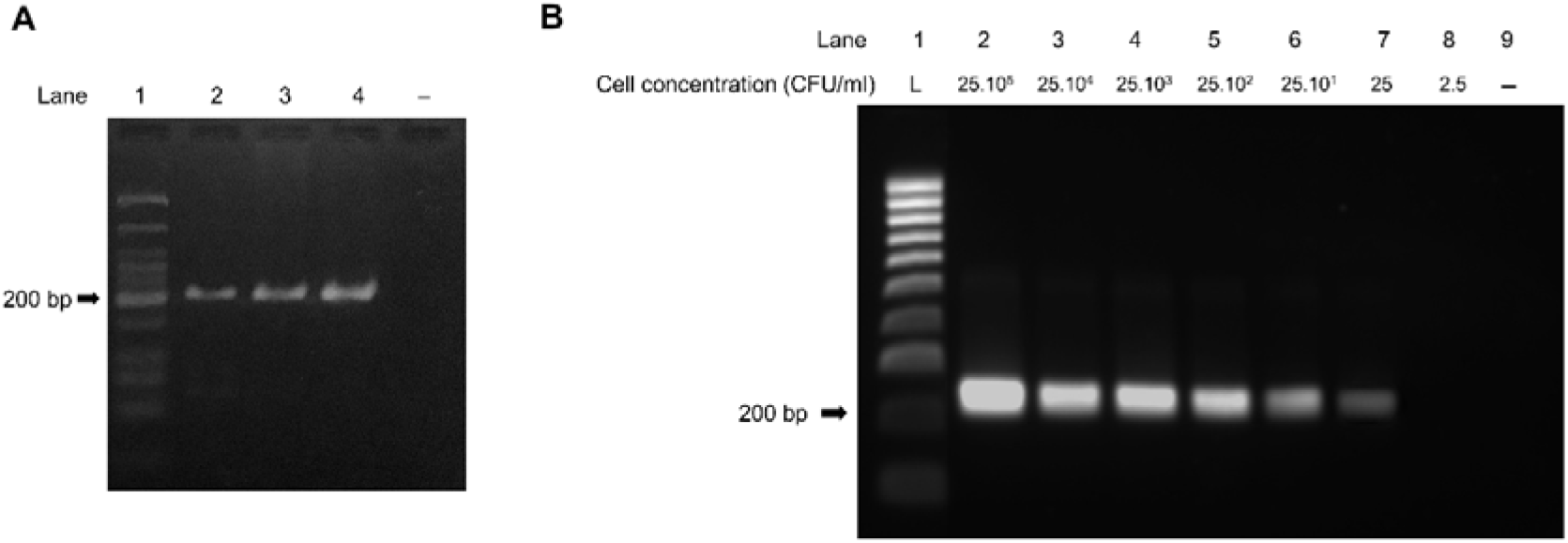
Direct RPA reaction with *L. monocytogenes* cell culture without DNA extraction. (A) The RPA assays were performed with 1 μl of the cell culture (lanes 2 to 4). Sterile water was used as the negative control sample. (B) Evaluating the LOD of direct *L. monocytogenes* RPA using the ten-fold serial dilution of the cell culture. Abbreviation, L: DNA ladder.

### Direct detection of *L. monocytogenes* cells by RPA in contaminated milk

To examine the ability of direct RPA assay using food sample, the artificially contaminated milk was prepared by spiked with *L. monocytogenes* cells at low concentrations ranging from 2.5 × 10^2^ to 2.5× 10^0^ CFU/ml. Without the need for DNA extraction or sample processed or cell enrichment, the LOD of direct RPA assay using milk samples was defined at 2.5 × 10^2^ CFU/ml (Fig. 6). The LOD value was ten times higher than that of direct RPA using cell culture, indicating that there are certain components in the milk sample interfering with the RPA reaction to some extent.

**Figure 6.**
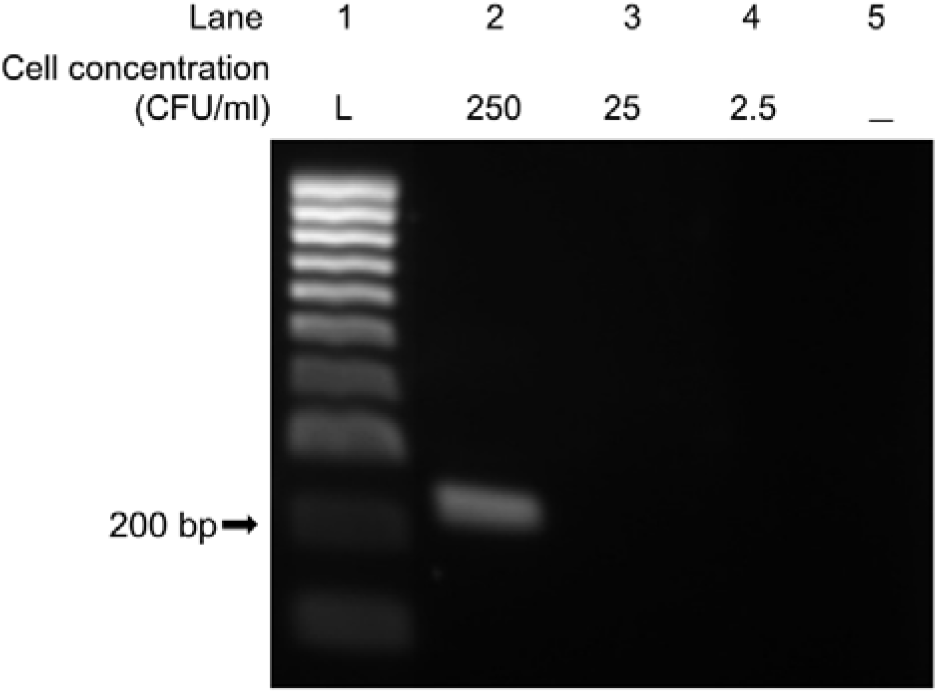
Performance of RPA assays for direct detection of *L. monocytogenes* in artificially contaminated milk. The RPA reactions were examined using 1 μl of the simulated infection milk spiked with different concentrations of *L. monocytogenes* cells ranging from 2.5 × 10^2^ to 2.5 × 10^0^ CFU/ml. Abbreviation, L: DNA ladder.

## DISCUSSION

*L. monocytogenes* is one of the common types of food poisoning bacteria which can cause serious foodborne diseases or even lethality. To promptly prevent the serious consequences of *L. monocytogenes* infection, the development of a rapid and efficient method for *L. monocytogenes* detection is needed. Currently, RPA is one of the isothermal amplification techniques that have been applied to detect different infectious bacteria precisely and quickly (**Liu *et al.*, 2017; Frimpong *et al.*, 2019; Geng *et al.*, 2019; Hu *et al.*, 2020**). In this study, the RPA assay was developed to specifically detect *L. monocytogenes* in direct crude samples.

The RPA amplification efficiency depends on the target sequence, amplicon size, and quality and type of sample tested (**Daher *et al.*, 2016**). Most previous studies analyzed the RPA performance using genomic DNA extracted from enrichment solution or using the spiked-*L. monocytogenes* food samples that were boiled or pretreated with lysis buffer to release the DNA (**Gao *et al.*, 2017; Du *et al.*, 2018**). These steps made the RPA assays previously developed more time-consuming, limiting their application at the field. In this study, we eliminated the genomic DNA extraction and used contaminated milk directly for the RPA reaction. The approach makes the testing handling simpler and faster in the diagnosis of *L. monocytogenes*. The *L. monocytogenes* RPA assay is advantageous due to no requirement for a specialized thermocycler. The assay could efficiently amplify the target sequence within 25 min at a low temperature of 39°C. The short testing time and low incubation temperature are beneficial for the early detection of *L. monocytogenes* in practical application. These advantages make the RPA assay developed to detect *L. monocytogenes* time-saving and cost-effective at the field with restricted resources.

The developed RPA assay had high specificity and sensitivity. No cross-reactivity was observed with several bacteria tested, which is in agreement with the previous study. Besides, the method has been proven that it was extremely sensitive compared to the PCR assay in this study. The LOD of the RPA assay was 310 fg of extracted DNA, indicating a 1000-fold higher sensitivity than PCR. Also, the RPA assay could directly detect as low as 25 CFU/ml of *L. monocytogenes* cells in medium cultures and 2.5 × 10^2^ CFU/ml of *L. monocytogenes* cells in contaminated milk. The obtained results agree with the previous studies, showing that RPA assays succeeded to directly detect the target bacteria in simulated clinical samples without the need for genomic DNA extraction (**Santiago-Felipe *et al.*, 2015; Geng *et al.*, 2019**). The reduced sensitivity of RPA observed with milk samples is mostly due to the matrices of crude samples that potentially affect the amplification process (**Miao *et al.*, 2019; L. Wang *et al.*, 2020**).

In summary, the direct RPA assay developed is a specific and rapid approach to alternate the traditional methods for efficiently and accurately diagnosing *L. monocytogenes* in food. Further evaluation of the assay with different types of crude samples by clinical testing for the diagnosis of *L. monocytogenes* infection, particularly in areas with restricted settings should be performed.

## CONCLUSION

The direct RPA assay which is rapid and sensitive developed in this study could be an alternative method for the diagnosis of *L. monocytogenes* infection, especially in areas with limited resources.

## Acknowledgments

This word was funded by NTTU Foundation for Science and Technology Development under grant number 2020.01.015.

## Notes

### Competing Interest Statement

The authors have declared no competing interest.

